# Detailed dendritic excitatory/inhibitory balance through heterosynaptic spike-timing-dependent plasticity

**DOI:** 10.1101/056093

**Authors:** Naoki Hiratani, Tomoki Fukai

**Affiliations:** Laboratory for Neural Circuit Theory, RIKEN Brain Science Institute, Hirosawa 2-1, Wako, Saitama, Japan 351-0198; Current address: Gatsby Computational Neuroscience Unit, University College London, 25 Howland Street, London, United Kingdom W1T 4JG

## Abstract

Balance between excitatory and inhibitory inputs is a key feature of cortical dynamics. Such balance is arguably preserved in dendritic branches, yet its underlying mechanism and functional roles remain unknown. Here, by considering computational models of heterosynaptic spike-timing-dependent plasticity (STDP), we show that the detailed excitatory/inhibitory balance on dendritic branch is robustly achieved through heterosynaptic interaction between excitatory and inhibitory synapses. The model well reproduces experimental results on heterosynaptic STDP, and provides analytical insights. Furthermore, heterosynaptic STDP explains how maturation of inhibitory neurons modulates selectivity of excitatory neurons in critical period plasticity of binocular matching. Our results propose heterosynaptic STDP as a critical factor in synaptic organization and resultant dendritic computation.

**Significance statement:** Recent experimental studies have revealed that relative spike timings among neighboring Glutamatergic and GABAergic synapses on a dendritic branch significantly influences changes in synaptic efficiency of these synapses. This heterosynaptic form of spike-timing-dependent plasticity (STDP) is potentially important for shaping the synaptic organization and computation of neurons, but its functional role remains elusive. Here, through computational modeling, we show that heterosynaptic plasticity causes the detailed balance between excitatory and inhibitory inputs on the dendrite, at the parameter regime where previous experimental results are well reproduced. Our result reveals a potential principle of GABA-driven neural circuit formation.

## Introduction

Activity dependent synaptic plasticity is essential for learning. Especially, spike time difference between presynaptic and postsynaptic neurons is a crucial factor for synaptic learning (Bi and Poo, 1998)(Caporale and Dan, 2008). Recent experimental results further revealed that the relative spike timings among neighboring synapses on a dendritic branch have significant influence on changes in synaptic efficiency of these synapses (Tsukada et al., 2005)(Hayama et al., 2013)(Paille et al., 2013)(Oh et al., 2015)(Bazelot et al., 2015). Especially, the timing of GABAergic input exerts a great impact on synaptic plasticity at nearby glutamatergic synapses. Similar phenomenon were also observed in biophysical simulations (Cutsuridis, 2011)(Bar-Ilan et al., 2013). This heterosynaptic form of spike-timing-dependent plasticity (h-STDP) is potentially important for synaptic organization on dendritic tree, and resultant dendritic computation (Mel and Schiller, 2004)(Branco et al., 2010). However, the functional role of h-STDP remains elusive, partly due to lack of simple analytical model.

In the understanding of homosynaptic STDP, simple mathematical formulation of plasticity has been playing important roles (Gerstner et al., 1996)(Song et al., 2000)(Vogels et al., 2011). Motivated by these studies, we constructed a mathematical model of h-STDP based on calcium-based synaptic plasticity models (Shouval et al., 2002)(Graupner and Brunel, 2012), and then considered potential functional merits of the plasticity. The model reproduces several effects of hSTDP observed in the hippocampal CA1 area and the striatum of rodents (Hayama et al., 2013)(Paille et al., 2013), and provides analytical insights for the underlying mechanism. The model further indicates that hSTDP causes the detailed balance between excitatory and inhibitory inputs on a dendritic branch owing to the inhibitory inputs that shunt long-term depression (LTD) at neighboring correlated excitatory synapses. This result suggests that not only the number and the total current of excitatory/inhibitory synapses are balanced at a branch (Liu, 2004)(Wilson et al., 2007), but temporal input structure is also balanced as observed in the soma (Dorrn et al., 2010)(Froemke, 2015). Moreover, by considering dendritic computation, we show that such detailed balance is beneficial for detecting changes in input activity. The model also reconciles with critical period plasticity of binocular matching observed in V1 of mice (Wang et al., 2010)(Wang et al., 2013), and provides a candidate explanation on how GABA-maturation modulates the selectivity of excitatory neurons during development.

## Methods

In this study, we first constructed a model of dendritic spine, then based on that model, built models of a dendritic branch, and a dendritic neuron. We also created an analytically tractable model of a spine, by reducing the original spine model.

### Spine model: Calcium-based STDP model with current-based heterosynaptic interaction

Let us first consider membrane dynamics of a dendritic spine. Membrane potential of a spine is mainly driven by presynaptic inputs through AMPA/NMDA receptors, backpropagation of postsynaptic spike, leaky currents, and current influx/outflux caused by excitatory/inhibitory synaptic inputs at nearby synapses. Hence, we modeled membrane dynamics of spine *i* with the following differential equation:

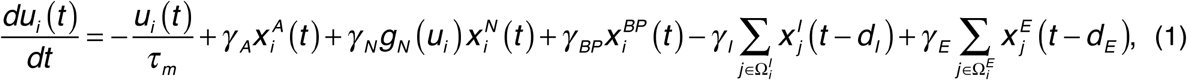

where *u_i_* is the membrane potential of the spine, and *τ_m_* is the membrane time constant(see Table 1 for definitions of variables). Here, conductance changes were approximated by current changes. The resting potential was renormalized to zero for simplicity. In next terms, 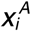 and 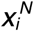 are glutamate concentration on AMPA/NMDA receptors respectively. The function *g_N_(u_i_)=α_N_u_i_+β_N_* represents voltage dependence of current influx through NMDA receptors. This positive feedback is enhanced when additional current is provided through back-propagation. As a result, the model reproduces large depolarization caused by coincident spikes between presynaptic and postsynaptic neurons. Although AMPA receptor also shows voltage dependence, here we neglected the dependence, as the relative change is small around the resting potential (Lüscher and Malenka, 2012). 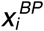 is the effect of backpropagation from soma, and the last two terms of the equation represents heterosynaptic current, which is given as the sum of inhibitory (excitatory) currents 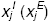 at nearby synapses. We defined sets of nearby inhibitory/excitatory synapses as 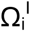 and 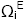 respectively, and their delays were denoted as *d_I_* and *d_E_*. Each input 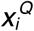 (Q = A,N,BP,I,E) is given as convoluted spikes:

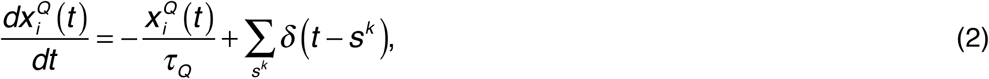

where *S^k^* represents the spike timing of the *k*-th spike. In the simulation, although convolution is calculated at the heterosynaptic synapse, this does not influence results because exponential decay is linear.

**Table 1.** Definitions of variables.

|  |  |  |
| --- | --- | --- |
| $u_i(t)$ | Membrane potential at spine $i$ | Eq. 1 |
| $c_i(t)$ | Calcium concentration at spine $i$ | Eq. 3 |
| $y_i(t)$ | Intermediate factor (interim synaptic weight) | Eq. 4 |
| $w_i(t)$ | Synaptic weight of spine $i$ | Eq. 5 |
| $g_N(u)$ | Voltage dependence of NMDA receptor | $g_N(u_i) = \alpha_N u_i + \beta_N$ |
| $g_V(u)$ | Voltage dependence of VDCC | $g_V(u_i) = \alpha_V u_i$ |
| $x_i^A(t)$ | Inputs through AMPA receptor | Eq. 2 (Q=A) |
| $x_i^N(t)$ | Inputs through NMDA receptor | Eq. 2 (Q=N) |
| $x_i^{BP}(t)$ | Back propagation | Eq. 2 (Q=BP) |
| $x_i^E(t)$ | Excitatory heterosynaptic inputs | Eq. 2 (Q=E) |
| $x_i^I(t)$ | Inhibitory heterosynaptic inputs | Eq. 2 (Q=I) |
| $u_b^k(t)$ | Membrane potential at dendritic branch $k$ | $u_b^k(t) = \sum_{i=1}^{N_b^E} w_i^k u_i^k(t) / (w_o^E N_b^E)$ |
| $u_{soma}(t)$ | Membrane potential at the soma | $u_{soma}(t) = \sum_k g_b(u_b^k(t))$ |
| $g_b(u)$ | Dendritic nonlinearity function | $g_b(u) = \begin{cases} u & (\text{if } u > u_b^o) \\ u_b^o & (\text{otherwise}) \end{cases}$ |

We next consider calcium influx to a spine through NMDA receptors and VDCC. For a given membrane potential *u_i_*, calcium concentration at spine *i* can be written as

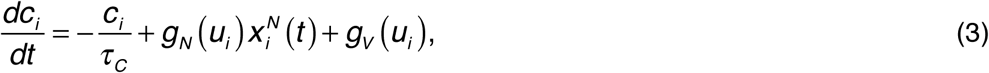

where *g_v_(u_i_)= α_v_u_i_* represents calcium influx through VDCC, and 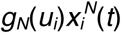 is the influx from NMDA.

Calcium concentration at spine is the major indicator of synaptic plasticity, and many results indicate that high Ca^2+^ concentration on a spine typically induces LTP, while low concentration often causes LTD (Lüscher and Malenka, 2012). Previous modeling studies revealed calcium-based synaptic plasticity model constructed on that principle well replicate various homosynaptic STDP time window observed in *in vitro* experiments (Shouval et al., 2002)(Graupner and Brunel, 2012). Hence, here we employed their framework for plasticity model. We additionally introduced an intermediate variable to reflect all-or-none nature of synaptic weight change (Petersen et al., 1998). This variable approximately represents the concentration of plasticity related enzymes such as CaMKII or PP1 (Graupner and Brunel, 2007). In the proposed model the intermediate *y_i_* and synaptic weight *w_i_* follow

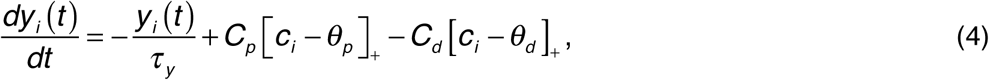

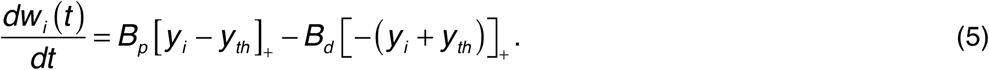

[X]_+_ is a sign function which returns 1 if X ≥ 0, returns 0 otherwise. Note that, in this model setting, as observed in recent experiments (Gambino et al., 2014), back-propagation is not necessary for LTP, if presynaptic inputs are given when the membrane potential at the spine is well depolarized.

In the simulation, we set common parameters as *τ_C_*=18.0ms, *τ_M_*=3.0ms, *τ_N_*=15.0ms, *τ_A_*=3.0ms, *τ_BP_*=3.0ms, *τ_I_*=3.0ms, *τ_E_*=6.0ms, *τ_Y_*=50s, *d_I_*=0.0ms, *α_N_*=1.0, *β_N_*=0.0, *α_V_* =2.0, *γ_A_*=1.0, *θ_p_*=70, *θ_d_*=35, *C_d_*=1.0, *B_p_*=0.001, *B_d_*=0.0005. Note that due to positive feedback between equations (1) and (3), effective timescales of calcium dynamics and NMDA channel become longer than the given values. In the model of STDP at striatum, in addition, we used *γ_N_*=0.05, *γ_BP_*=8.0, *γ_I_*=5.0, *C_p_*=2.3, *y_th_*=250, while for the model of Schaffer collateral synapses, we used *γ_N_*=0.2, *γ_BP_*=8.5, *γ_I_*=3.0, *C_p_*=2.2, *y_th_*=750, *d_E_*=1.0, *γ_E_*=1.0. In the parameter search, decay time constants were chosen from biologically reasonable ranges (Koch, 1998), *α_N_*, *γ_A_*, *С_d_*, *В_d_* were fixed at unitary values, and other parameters were manually tuned. Subsequently, robustness of parameter choice was confirmed numerically (Fig. 3). Synaptic weight variables {*w*} were bounded to 0 < *w* < 500, and initialized at *w* = 100. All other variables were initialized at zero in the simulation. Paired stimulation was given every 1 second for 100 seconds, and synaptic weight changes were calculated from the values 400 seconds after the end of stimulation. In the cortico-striatal synapse model, the inhibitory spike was presented at the same timing with the presynaptic spike, and for Schaffer collateral synapses, inhibitory spikes were given 10 milliseconds before pre (post) spikes in pre-post (post-pre) stimulation protocols. In calculation of intermediate variable *y(t)* in Figures 2B,D, we ignored the effect of exponential decay, because of the difference in timescale (*τ_y_* ≫ 1 seconds). We subtracted 7.5 milliseconds of axonal delay from the timing of presynaptic stimulation in the calculation of spike timing difference.

**Figure 1.**
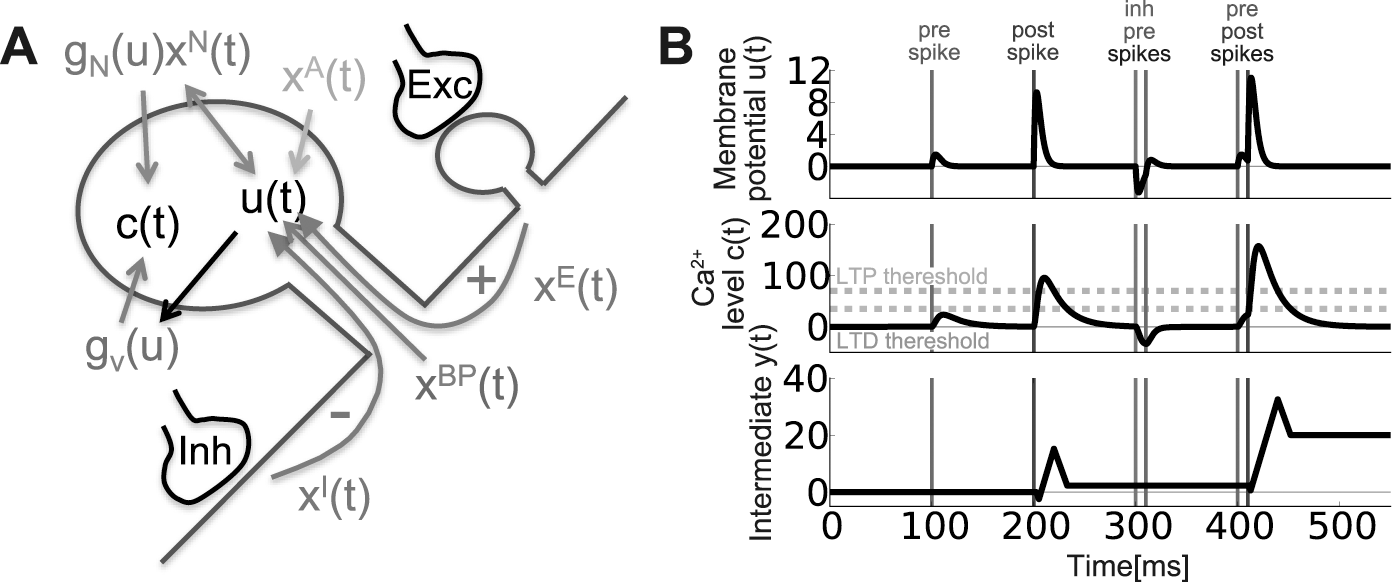
Schematic figure of the model of heterosynaptic spike-timing-dependent plasticity (h-STDP). **A)** A schematic figure of the model. Two variables in the spine *u(t)* and *c(t)* represent the normalized membrane potential and Ca^2+^ concentration respectively. Presynaptic action potentials modulate the membrane potential *u(t)* through AMPA (*X(_A_)*) and NMDA (*g_N_(u)x^N^*) receptors. In addition, *u(t)* is modified by back-propagation (*X(_BP_)*), and heterosynaptic current caused by excitatory (*X(_E_)*) and inhibitory (*X(_I_)*) inputs. Calcium level *c(t)* is modulated by influx/outflux through NMDA (*g_N_(u)x^N^*) and VDCC (*g_V_(u)*). Consequently, *c(t)* is indirectly controlled by *u(t)* because both NMDA and VDCC are voltage-dependent. **B)** An example of dynamics of the membrane potential variable *u(t)* (top), Ca^2+^ concentration *c(t)* (middle), and the intermediate variable *y(t)* that controls the synaptic weight *w(t)*(bottom). Change in the Ca^2+^ level roughly follows the membrane potential dynamics, and the intermediate variable *y(t)* is positively (negatively) modulated when Ca^2+^ level is above LTP(LTD) thresholds represented by orange(cyan) dotted lines. Based on the intermediate variable *y(t)*, synaptic weight *w(t)* is updated in a slow timescale (see Figure 5C for example).

**Figure 2.**
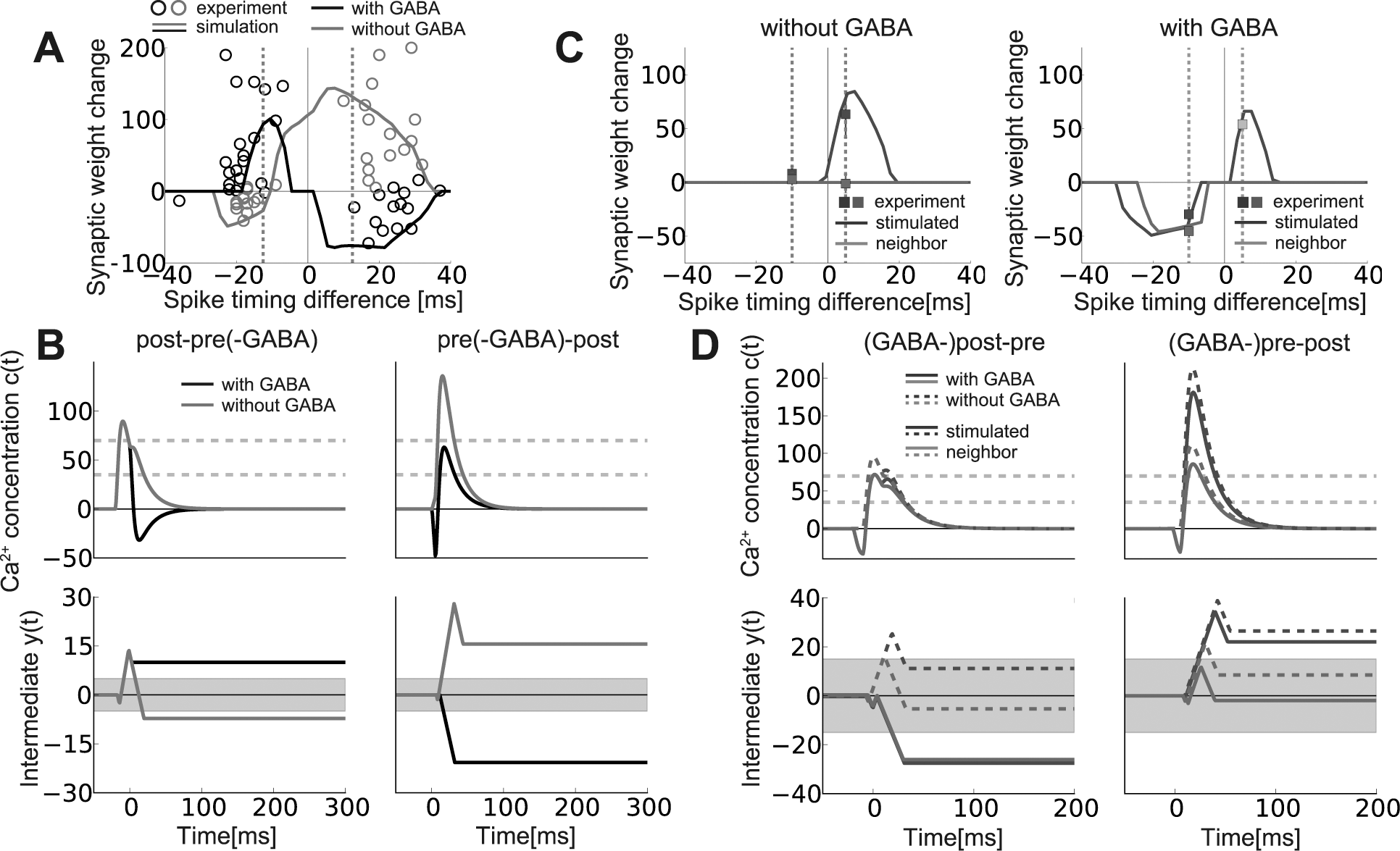
The model reproduces spike-timing-dependent heterosynaptic effects. **A)** Spike timing window with/without a di-synaptic GABAergic input. Lines are simulation data, and points are experimental data taken from (Paille et al., 2013). Vertical dotted lines represent the spike-timing differences at which Panel **B** was calculated. **B)** Dynamics of calcium concentration *c(t)* (top) and the intermediate variable *y(t)* (bottom) at the stimulated spine. Gray areas in the bottom figures represent the regions satisfying *y(t)* < *y(t)*/*K_rep_*, in which the change in the intermediate is not reflected into synaptic weight, where *K_rep_* represents the number of paired stimulation given in the simulation for Panel **A**. **C**) Synaptic weight change with/without GABAergic inputs right before pre/post stimulation. Data points were taken from (Hayama et al., 2013). The cyan point is a result from muscimol application, not GABA uncaging. **D)** Dynamics of *c(t)* and *y(t)* at the stimulated spine (blue lines) and a neighboring spine (green lines). Solid lines represent the dynamics under GABA uncaging, and dotted lines are the controls. Note that in the left panel, fractions of blue lines are hidden under green lines, because postsynaptic and inhibitory heterosynaptic inputs cause the same dynamics in both stimulated and neighboring spines.

**Figure 3.**
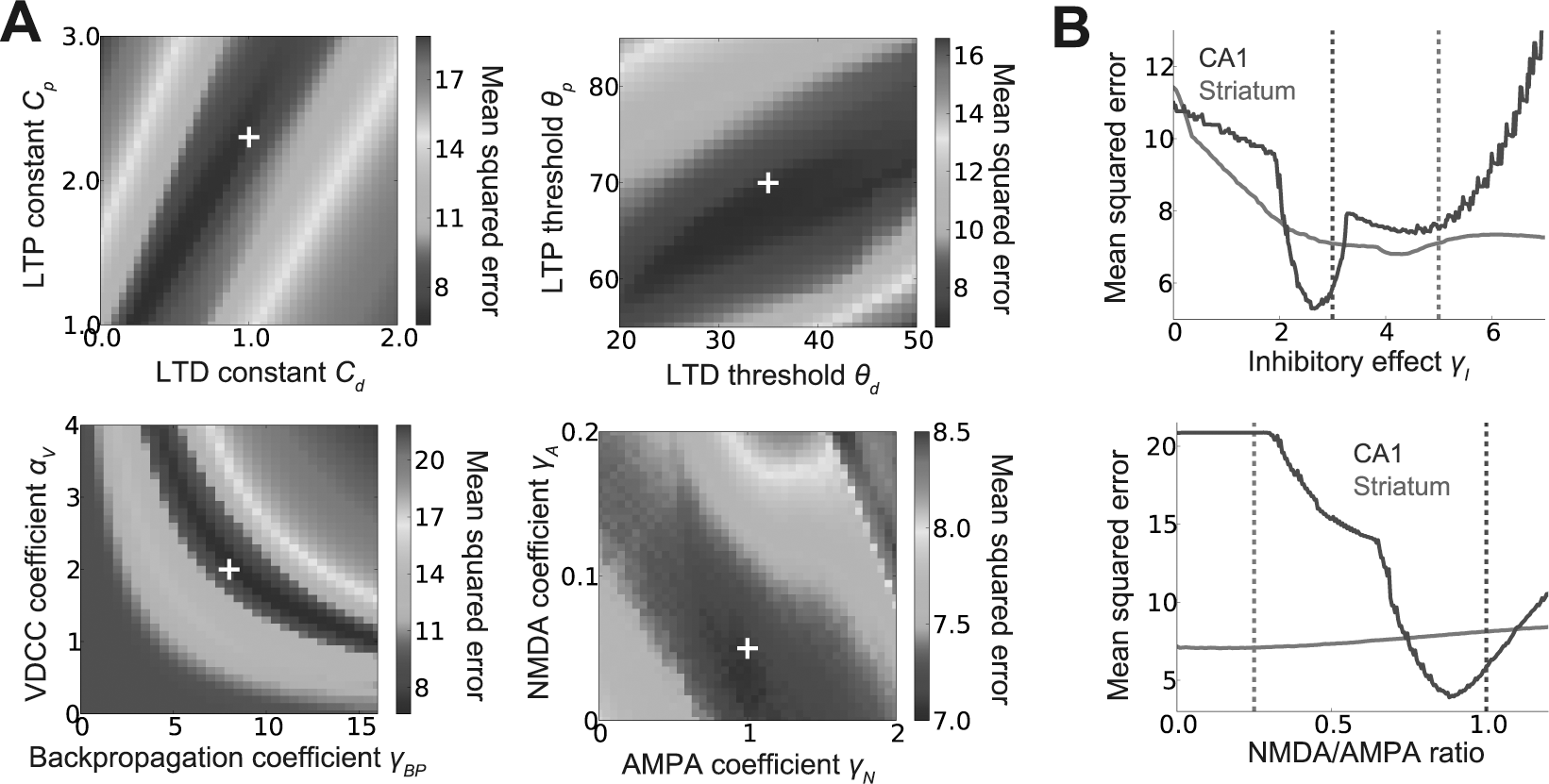
Parametric robustness of h-STDP model. **A)** Mean squared fitting errors for the model of the striatum experiment at various values of model parameters. In each panel, all other parameters were fixed, and the white mark represents the value used in Figure 2. See Methods for the definitions of parameters. **B)** Comparison of parametric dependence of models fitted for the results from the striatum and CA1 experiments. Vertical dotted lines represent the value used in Figure 2. NMDA/AMPA ratio in the bottom panel was calculated as *γ_N_τ_N_*/*γ_A_τ_A_* at various values of *γ_N_*. Note that the fluctuation in blue lines was caused by double-threshold dynamics of the model, not by noise.

### Dendritic hotspot model

Dendritic hotspot model was constructed based on the Schaffer collateral synapse model described above. For simplicity, we hypothesized that heterosynaptic current due to inhibitory spike arrives on excitatory spines at the same time, and also disregarded E-to-E interaction by setting *γ_E_*=0.0. Correlated spikes were generated using hidden variables as in previous studies (Vogels et al., 2011)(Hiratani and Fukai, 2015). We generated five dynamic hidden variables, and updated them at each time step by 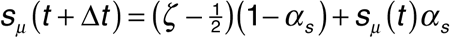, where *α_s_* = exp[−Δ*t*/*τ_S_*], *τ_S_*=10ms, *μ*=0, 1,…,4, and *ζ* is a random variable uniformly chosen from [0,1). In the simulation, the time step was set at Δ*t*=0.1ms. Activities of presynaptic neurons were generated by rate-modulated Poisson process with 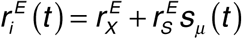 excitatory neuron *i* modulated by the hidden variable *μ* (due to non-negative constraint on 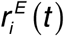, we set 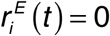 when 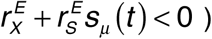. Similarly, the presynaptic inhibitory neuron was described by a Poisson-model with 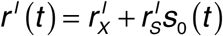. Activity of the postsynaptic neuron was given as a Poisson-model with a fixed rate *r_post_*. We set parameters 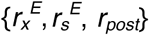 in a way that all pre and postsynaptic excitatory neurons show the same average firing rate 157 at 5Hz, to avoid the effect of firing-rate difference on synaptic plasticity.

For parameters, we used *γ_I_*=1.2, *β_N_*=1.0, *γ_BP_*=8.0, *C_p_*=2.11, *y_th_*=250 and other parameters 159 were kept at the same value with the original Schaffer collateral model. Except for Figure 5D, the delay of inhibitory spike was set as zero. Presynaptic activities were given by 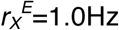, 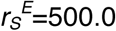, 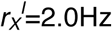, 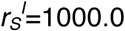, and postsynaptic firing rate was set as *r_post_* = 5.0Hz. In Figure 5C, the correlation was calculated between dendritic membrane potential *g_b_(u_b_)* and hidden variables {*s_μ_(t)*}, where 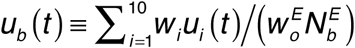, and *g_b_*(*u*) was defined as *g_b_*(*u*)=*u* if 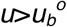, otherwise 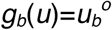 with 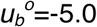.

**Figure 4.**
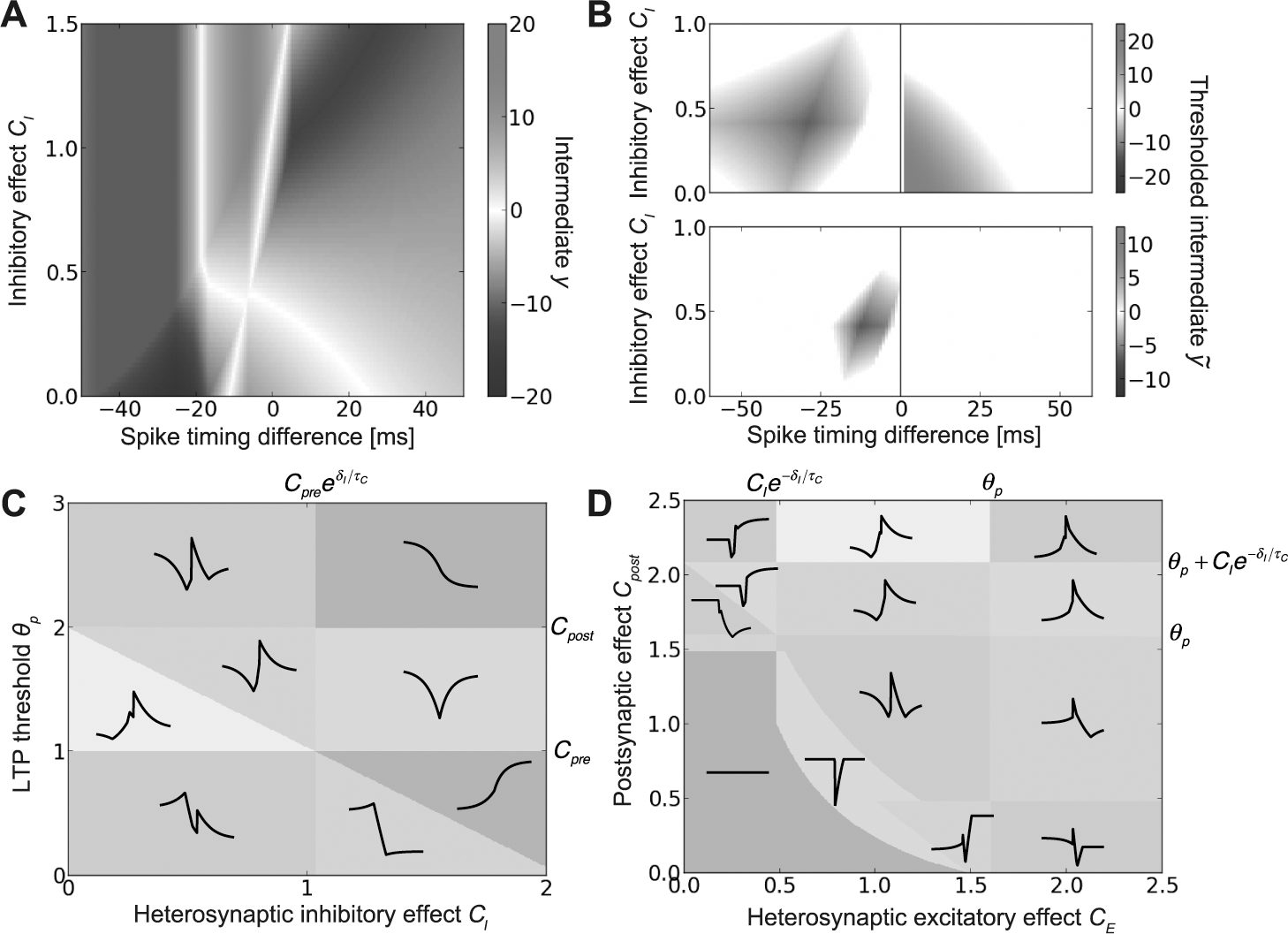
Phase transitions on STDP time window in an analytical model of h-STDP. **A, B)** STDP windows at various strength of heterosynaptic inhibitory effect *C_I_*. Panel **A** corresponds to the striatum experiment, and Panel **B** corresponds to the CA1 experiment. Top and bottom figures in Panel **B** represent the stimulated and a neighbor spine, respectively. Note that values in Panel **B** were calculated by *ỹ* = *sgn*(*y*)·[|*y*|−15]_+_ to reflect the effect of thresholding. **C)** Phase diagram of STDP time window calculated for inhibitory effect *C_I_* and LTP threshold *θ_p_*. Colors show the number of local minimum/maximum, and lines are typical STDP time windows at each phase. Parameters written on the right side (top) of the panel represent the critical values of *θ_p_* (*C_I_*). **D)** Phase diagram calculated for heterosynaptic excitatory effect parameter *C_E_* and postsynaptic effect parameters *C_post_* at a fixed inhibitory effect (*C_I_*=0.5). See Reduced model in Methods for details.

**Figure 5.**
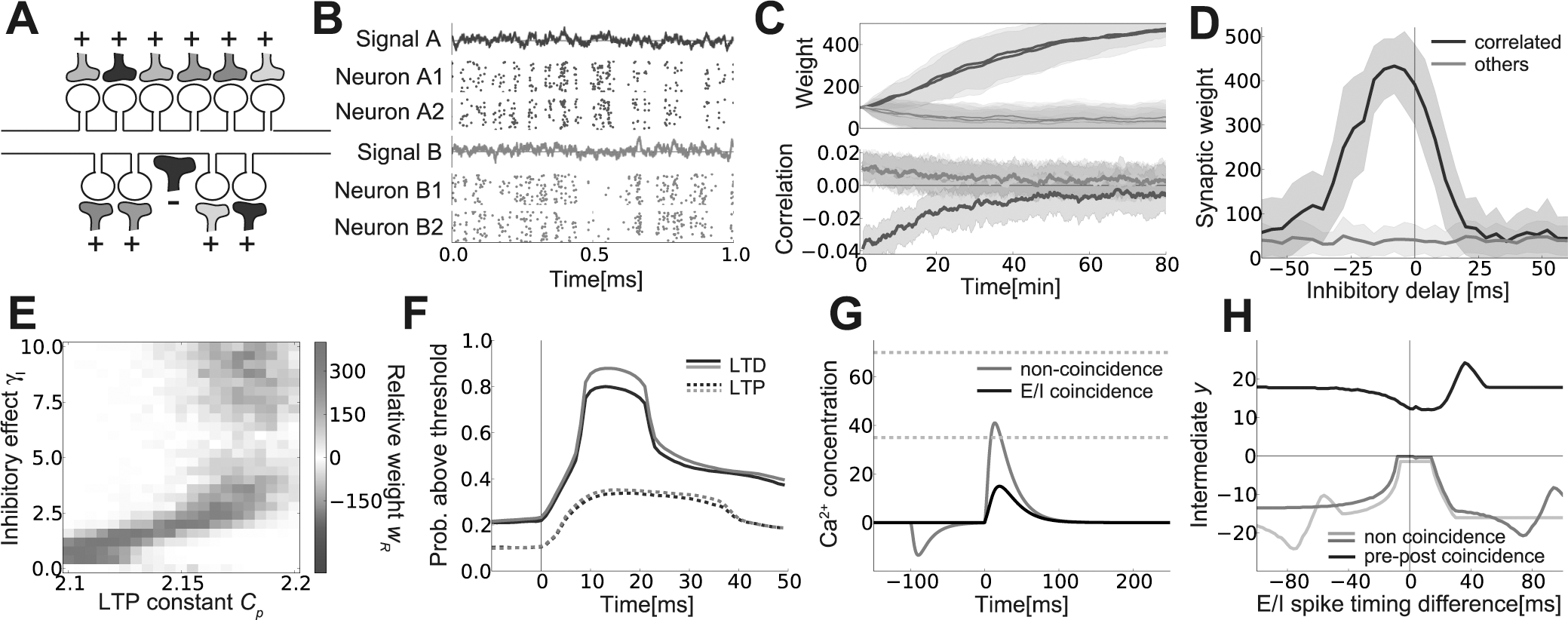
Emergence of detailed dendritic excitatory/inhibitory balance by h-STDP. **A)** A schematic figure of a dendritic hotspot model. The shaft synapse represents an inhibitory input. Colors represent spike correlation between synaptic inputs. **B)** Examples of correlated spike inputs. Each Raster plot was calculated from 50 simulation trials. **C)** Changes in synaptic weight w (top) and the correlation between the dendritic membrane potential and hidden signals (bottom) under h-STDP. The blue lines represent dynamics of synapses correlated with the inhibitory input. **D)** Synaptic weight change at the excitatory synapses correlated with the inhibitory inputs (blue) and at other synapses (gray), at various inhibitory delays. Error bars in Panels **C** and **D** represent standard deviations over 50 simulation trials. **E)** Relative weight changes *w_R_* calculated at various parameters. We defined *w_R_* by 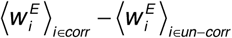 where "corr" represents a set of excitatory synapses correlated with the inhibitory synapse, and "un-corr" stands for uncorrelated ones. Here, weights were calculated by taking average over 10 simulations. **F)** Probability of LTP/LTD occurrence after a presynaptic spike calculated from a simulation. Lines represent the mean LTP/LTD probabilities at excitatory synapses correlated with the inhibitory input (blue lines) and other synapses (gray lines), respectively. **G**, **H**) Results in single-spike simulations. E/I coincidence prevents LTD effect due to pre-spike (**G**), without affecting LTP effect due to pre-post coincidence (**H**). In Panel **G**, inhibitory spikes were provided at *t*=0 in the black line, *t*=-100ms in the gray line, and the excitatory presynaptic spike was given at *t*=0 in both lines. Similarly, in Panel **H**, postsynaptic spikes were provided at *t*=-75 (light-gray), 0 (black), +75ms (dark-gray), and the presynaptic spike was given at *t*=0 in all lines.

### Two-layered neuron model

Previous studies suggest that complicated dendritic computation can be approximated by a two-layered single cell model (Poirazi et al., 2003)(London and Häusser, 2005). Thus, we constructed a single cell model by assuming that each hotspot works as a subunit of a two-layered model. We defined the mean potential of a dendritic subunit *k* by 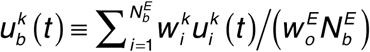, and calculated the somatic membrane potential by 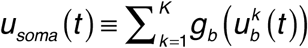. Postsynaptic spikes were given as a rate-modulated Poisson model with the rate *u_soma_*(t)/*I_dv_*(t). *I_dv_*(t) is the divisive inhibition term introduced to keep the output firing rate at *r_post_*. By using the mean somatic potential 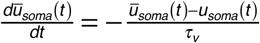, *I_dv_*(*t*) was calculated as *I_dv_* (*t*) ≡ *ū_soma_*(*t*)/*r_post_*.

In the simulations described in Figure 6, we used *C_p_*=2.01, *τ_v_*=1s, *K*=100, and other parameters were kept at the same values with the dendritic hotspot model. During the learning depicted in Figure 6BC, we used the same input configuration with the dendritic hotspot model. In Figure 6DE, The activity levels of hidden variables {*s_μ_(t)*} were kept at a constant value (*s_μ_(t)*=0.25) during 500ms stimulation, and otherwise kept at zero. Additionally, inhibitory presynaptic activities were set as 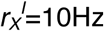, 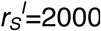. In Figure 6D, we modulated firing rates of both excitatory and inhibitory presynaptic neurons, by changing the activity levels of hidden variables {*s_μ_(t)*} from 0.1 to 0.5. The ratio of change detecting spikes was defined as the ratio of spikes occurred within 50 milliseconds from the change to the total spike count.

**Figure 6.**
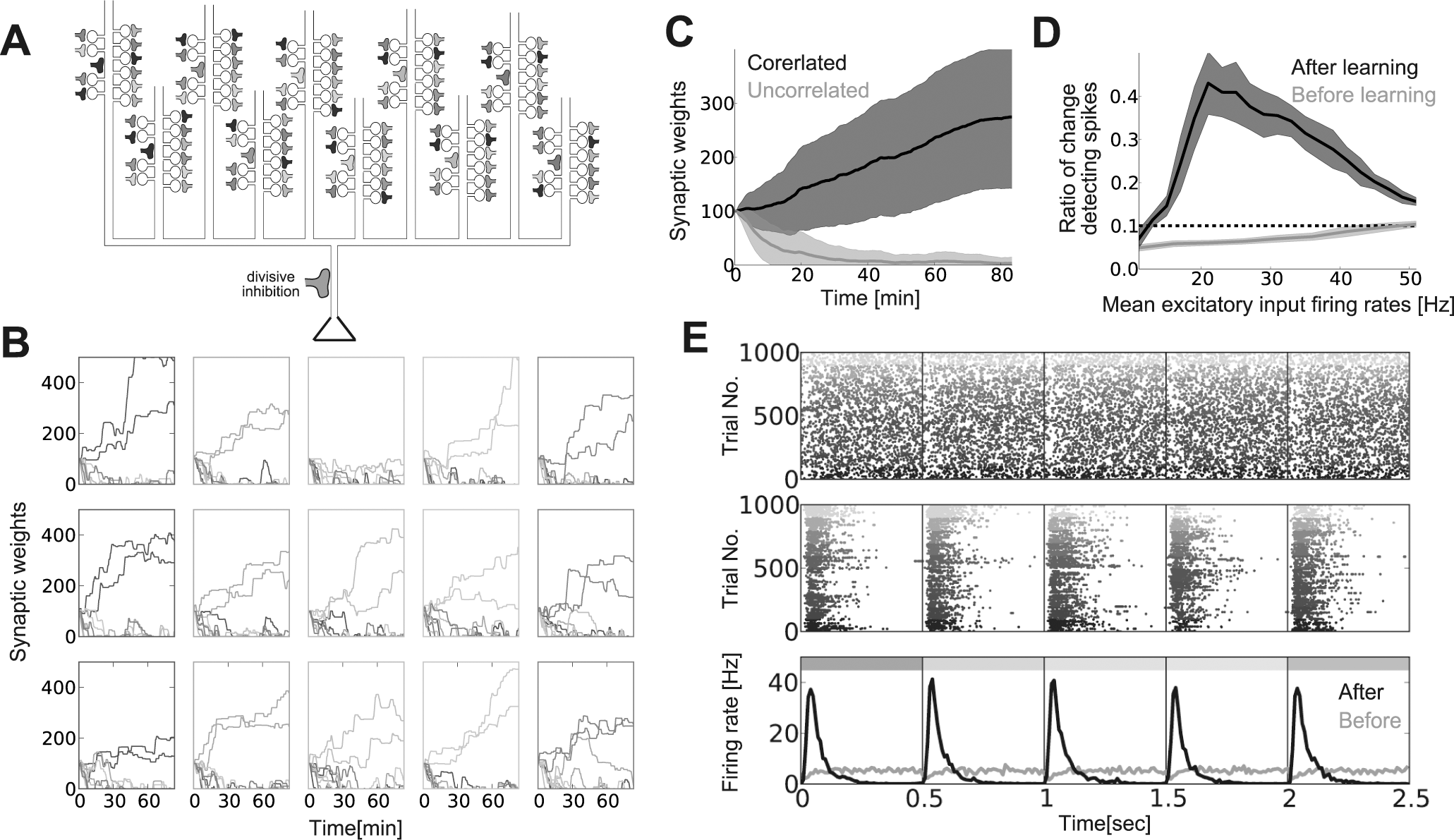
Detailed dendritic excitatory/inhibitory balance in a two-layered single cell model. **A)** A schematic illustration of the single cell model. The actual model has 100 dendritic branches each receiving 10 excitatory inputs and 1 inhibitory input. As in Figure 5A, inhibitory inputs are represented by shaft synapses. **B)** Examples of synaptic weight change at each branch. The color of the frames represents the selectivity of the inhibitory input to the branch. Each row represents a different simulation trial. **C)** Mean synaptic weight dynamics of synapses correlated to the local inhibitory inputs, and other synapses. **D)** The ratio of change detecting spikes before and after learning. The ratio was defined as the fraction of spikes occurs within 50 milliseconds from a change in stimuli to the total. In the x-axis, in addition to the mean excitatory input firing rates, the mean inhibitory input firing rates were also modulated from 50 Hz to 210Hz correspondingly to keep the E/I balance of the input. **E)** Raster plots of output spikes before (top) and after (middle) learning, and their firing rate dynamics (bottom), averaged over 100 trials each for 10 simulated neurons. Colors of spikes in the Raster plots represent results from different simulation trials. Black vertical lines represent the change points of excitatory inputs, and horizontal colored bars at the top of the bottom panel corresponds to the colors of presynaptic neurons active in each period. We introduced 10 milliseconds delay between excitatory and inhibitory stimulus both during learning (panel **B** and **C**) and in the change detecting task (panel **D** and **E**). The averages in panel **C** and **D** were taken over 10 simulation trials.

For the model of critical period plasticity of binocular matching depicted in Figure 7, we also used the two-layered single cell model. The neuron model has *K*=100 dendritic branches, each receives 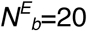 excitatory inputs and 1 inhibitory input. At each branch, half of excitatory inputs are from the contralateral eye, and the other half are from the ipsilateral eye. Each excitatory input neuron have direction selectivity characterized with 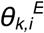, and shows rate-modulated Poisson firing with

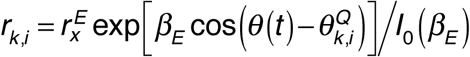

where *θ(t)* is the direction of the visual stimulus at time *t*, Q is either contra- or ipsilateral, and *I_0_*(*β_E_*) is the modified Bessel function of order 0. Similarly, firing rate of an inhibitory neuron was given as 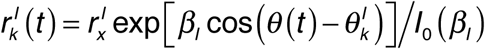. For each excitatory input neuron, mean direction selectivity 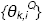 was randomly chosen from a von Mises distribution 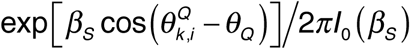, where Q={contra, ipsi}. In the simulation, we used *θ_contra_*=-π/4, *θ_ipsi_*=-π/4. Correspondingly, mean direction selectivity of a inhibitory neuron 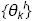 was defined as the mean of its selectivity for ipsi- and contralateral inputs (ie. 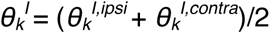), where 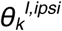 and 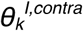 were also randomly depicted from 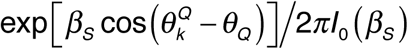. Direction of visual stimulus *θ(t)* changes randomly with *θ*(*t* + Δ*t*) = *θ* (*t*)+*σ_sr_ζ_G_* where *ζ_G_* is a Gaussian random variable, and *Δt* is the time step of the simulation. To mimic monocular deprivation, in the shadowed area of Figure 7E, we replaced contra-driven input neuron activity with a Poisson spiking with constant firing rate 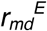. In addition, to simulate the lack of contra-driven inputs to inhibitory neurons, we replaced inhibitory activity with 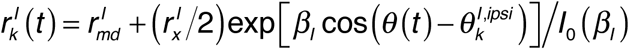. Similarly, in Figure 7C, we measured direction selectivity by providing monocular inputs, while replacing the inputs from the other eye with a homogeneous Poisson spikes with firing rate 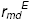.

**Figure 7.**
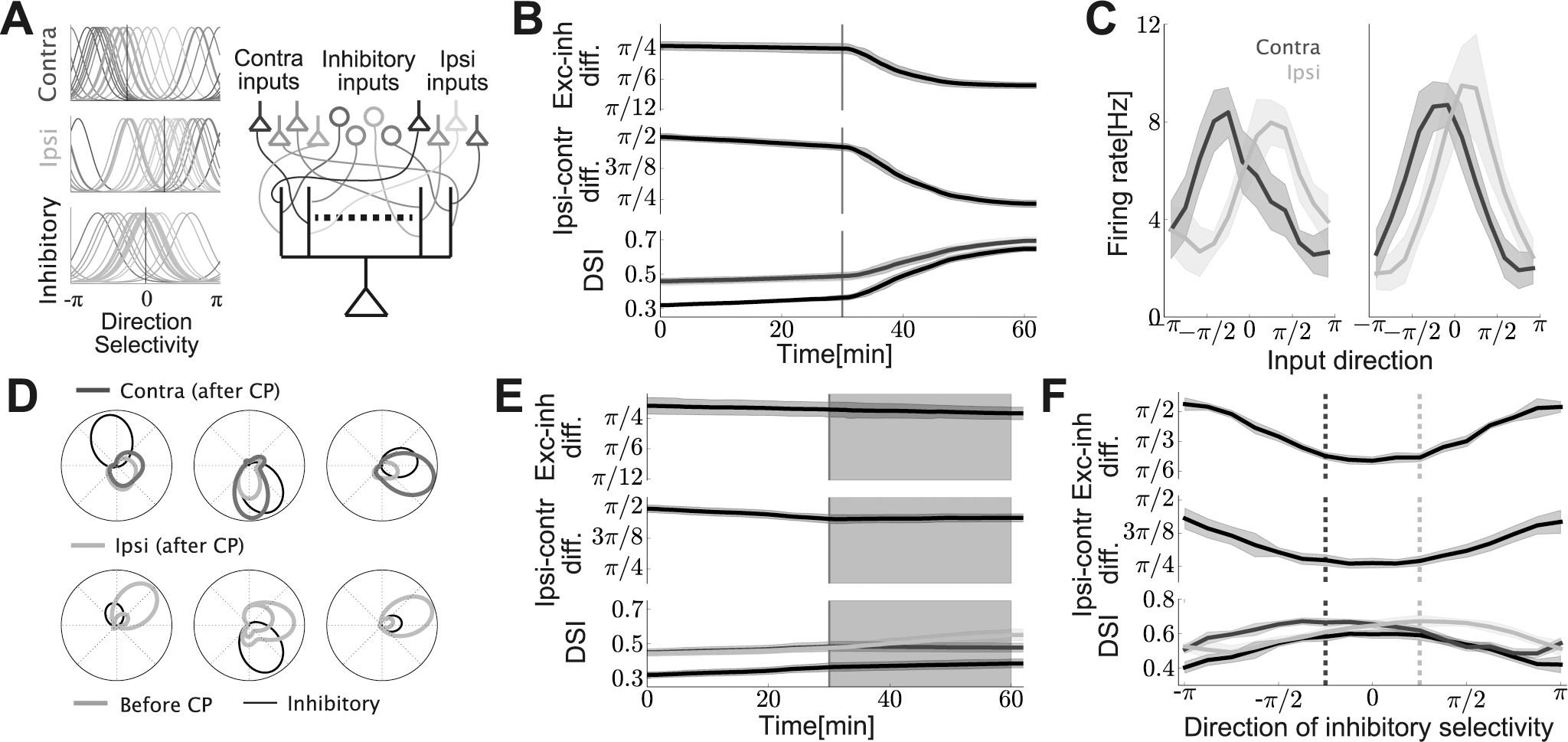
h-STDP can trigger binocular matching. **A)** (left): Direction selectivity of input neurons. In the model, as depicted by black vertical lines,majorities of excitatory input neurons from the contralateral (ipsilateral) eye are selective fordirections around *θ*=−*π* 4(*θ* = *π*/4), while inputs from the inhibitory neurons are weakly selective for *θ*=0. (right): A schematic figure of model configuration. Each dendritic branch receives inputs from both ipsi-and contralateral driven excitatory neurons and also from inhibitory neurons. **B)** (top): Difference between mean excitatory direction selectivity and inhibitory direction selectivity in each branch. (middle): Difference between mean ipsi-driven excitatory direction selectivity and mean contra-driven excitatory direction selectivity over all synapses on the neuron. (bottom): Direction selectivity index (DSI) calculated for contralateral inputs (purple), ipsilateral inputs (light-green; hidden under the purple line), and binocular inputs (black). See Neuron model in Methods for the details of evaluation methods. Red vertical lines represent the timing for introduction of inhibitory inputs. Throughout Figure 7, error bars are standard deviations over 10 simulation trials. **C)** Firing responses of the neuron for monocular inputs, right after the initiation of inhibitory inputs (left; *t*=30min), and after the learning (right; *t*=60min). **D)** Examples of direction selectivity of three representative branches before (gray lines; *t*=0min) and after (purple/light-green lines; *t*=60min) the learning. Black lines represent the selectivity of the inhibitory input to the branch. **E)** Behavior in monocular deprivation model. In shadowed areas, to mimic monocular deprivation, contra-driven inputs were replaced with rate-fixed Poisson inputs. Ordinates are the same with Panel **B. F**) Synaptic weights development at different mean inhibitory selectivity. Ordinates are the same with Panel B, and values were calculated at *t*=60min. Purple and green vertical dotted lines are mean selectivity of contra-and ipsilateral excitatory inputs, respectively.

To evaluate the development of binocular matching, we introduced three order parameters. First, the difference between mean excitatory direction selectivity and inhibitory selectivity at a *k* branch was evaluated by 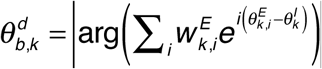. Similarly, the global direction selectivity difference between inputs from the ipsi- and contralateral eyes were defined by

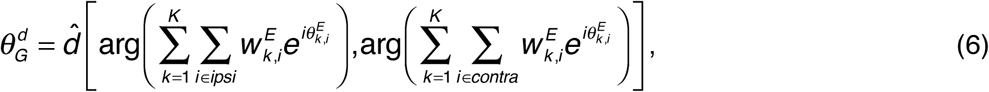

where the function 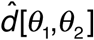 calculates the phase difference between two angles. Finally, direction selectivity index DSI for binocular input was calculated by

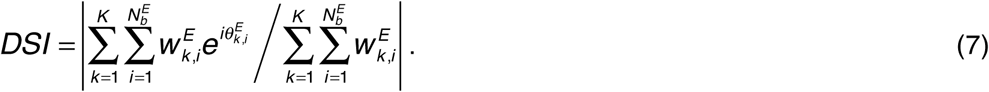

For the calculation of the monocular direction selectivity index, at each branch *k*, we took sum over 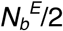 excitatory inputs corresponding to each eye instead of all 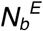 inputs.

In the simulation, we set *γ_I_*=2.5, *C_p_*=1.85, *y_th_*=750.0, 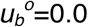, and the rest of parameters were kept at the values used in the dendritic hotspot model. Inputs parameters were set at *β_E_*=4.0, *β_I_*=2.0, *β_S_*=1.0, *θ_contra_*=-π/4, *θ_ipsi_*=-π/4, 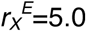, 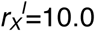, 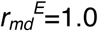, 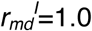, and 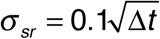.

### Reduced model

If we shrink equations for membrane potential (eq. 1) and calcium concentration (eq. 3) into one, the reduced equation would be written as,

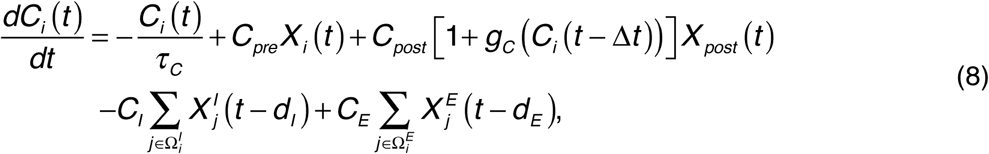

where *g_c_*(*X*) = [*X*]_+_ *ηX* captures the nonlinear effect caused by pre-post coincidence (i.e. (*g_c_*(*X*) returns *ηX* if *X*>0, otherwise returns 0). All inputs *X_i_*, *X_post_*, 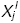, 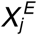 were given as point processes, and *d_I_*, *d_E_* are heterosynaptic delays. *g_c_* was calculated from the value of *C_i_* at *t*=*t*-*Δt* to avoid pathological divergence due to the point processes. In the simulation, we simply used value of *C_i_* one time step before. For the intermediate *y*, we used the same equation as before. Note that above equation is basically same with the one in (Graupner and Brunel, 2012) except for the nonlinear term *g_c_*(*C*) and the heterosynaptic terms.

Let us consider weight dynamics of an excitatory synapse that has only one inhibitory synapse in its neighbor. For analytical tractability, we consider the case when presynaptic, postsynaptic, and inhibitory neurons fire only one spikes at *t=t_pre_*, *t_post_*, *t_I_* respectively. In case of the CA1 experiment, because GABA uncaging was always performed before pre and postsynaptic spike, the timing of inhibitory spike is given as *t_I_* = min(*t_pre_*,*t_post_*)–*δ_I_* for *δ_I_* < 0. In this setting, the change in intermediate variable of the excitatory synapse is given as

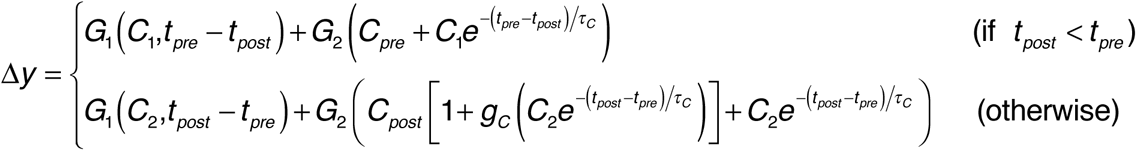

where,

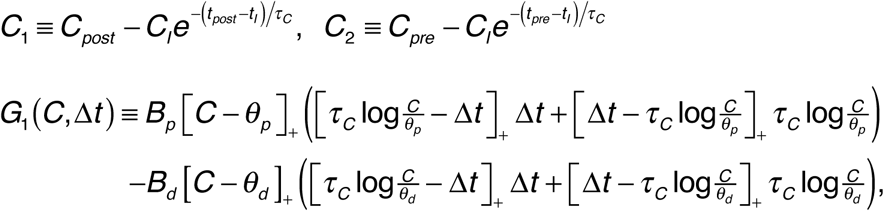

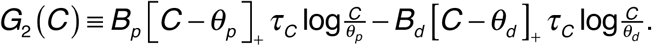

Similarly, in case of the striatum experiment, by setting *η*=0, the change in the intermediate variable is given as

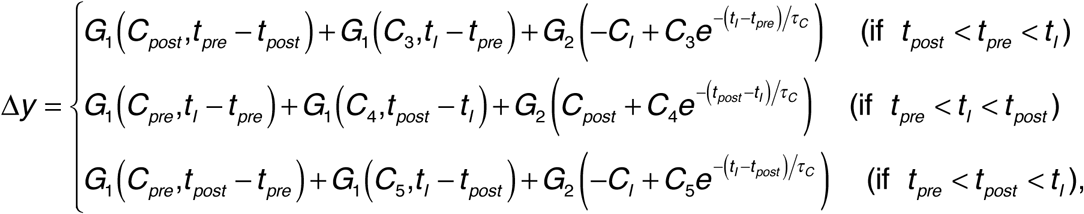

where

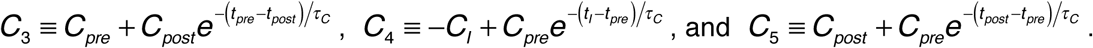

In the simulation, parameters were set at *τ_c_*=30ms, *C_post_*=2.0, *θ_p_*=1.6, *θ_d_*=1.0, *B_p_*=2.25, *B_d_*=1.0. Additionally, in the model of a Schaffer collateral synapse, we used *δ_I_*=1.0, *C_pre_*=1.0, *C_E_*=0.30, *η*=2.0, and for the model of a cortico-striatal synapse, we employed *δ_I_*=5.0, *C_pre_*=0.75, *C_E_*=0.0, *η*=0.0. In Figures 3C and D, we used the parameter set for the model of Schaffer collateral synapse.

As depicted in Figure 3D, the model also provides an analytical insight to E-to-E interaction, in addition to I-to-E interaction analyzed in the main result. In E-to-E interaction, neighboring synapses receive small heterosynaptic calcium transient *C_E_* instead of presynaptic input *C_pre_*. Thus, we can characterize the shapes of STDP time windows by the heterosynaptic excitatory effect parameter *C_E_*, and postsynaptic effect parameters *C_post_* (Fig. 3D). When the postsynaptic effect parameter *C_post_* satisfies *θ_p_* < *C_post_* < *θ_p_* + *C_I_e*^−*δ_I_*/*τ_c_*^, and the heterosynaptic effect parameter *C_E_* fulfills *C_I_e*^−*δ_I_*/*τ_c_*^ < *C_E_* < *θ_p_*, STDP time window shows Hebbian-type timing dependency (upper-middle orange-colored region in Fig. 3D). On the other hand, if *C_E_* is smaller than *C_I_e*^−*δ_I_*/*τ_c_*^ while satisfying *θ_p_* + *C_I_e*^−*δ_I_*/*τ_c_*^ – *C_post_* < *C_E_*, then the STDP curve becomes LTD dominant (upper-left green-colored region in Fig. 3D) as observed in experiments (Hayama et al., 2013)(Oh et al., 2015). Excitatory heterosynaptic effect *C_E_* is expectedly smaller than the inhibitory effect *C_I_*, because the inhibitory potential is typically more localized (Gidon and Segev, 2012). Thus, *C_E_* < *C_I_e*^−*δ_I_*/*τ_c_*^ is also expected to hold for small *δ_I_*, suggesting robust heterosynaptic LTD at neighboring synapses.

## Results

### Calcium-based synaptic plasticity model with current-based heterosynaptic interaction explains h-STDP

We constructed a model of a dendritic spine as shown in Figure 1A (see Spine model in Methods for details). In the model, the membrane potential of the spine *u(t)* is modulated by influx/outflux from AMPA/NMDA receptors (*x^A^* and *g_N_(u)x^N^* in Fig. 1A), back-propagation (*x^BP^*), and heterosynaptic currents from nearby excitatory/inhibitory synapses (*x^E^* and *x^I^*) (see Table 1 for the definitions of variables). Calcium concentration in the spine *c(t)* is controlled through NMDA receptors and voltage-dependent calcium channels (VDCC) *g_v_(u)* (Higley and Sabatini, 2012). Because both NMDA and VDCC are voltage-dependent (Lüscher and Malenka, 2012), the calcium level in the spine is indirectly controlled by pre, post, and heterosynaptic activities (Fig. 1B top and middle panels). For synaptic plasticity, we used calcium-based plasticity model, in which LTP/LTD are initiated if the Ca^2+^ level is above LTP/LTD thresholds (orange and cyan lines in Fig. 1B middle). This plasticity model is known to well capture homosynaptic STDP (Shouval et al., 2002)(Graupner and Brunel, 2012). We introduced an intermediate variable *y(t)* to capture non-graded nature of synaptic weight change (Petersen et al., 1998). Thus, changes in Ca^2+^ level are first embodied in the intermediate *y(t)* (Fig. 1B bottom), and then reflected to the synaptic weight *w(t)* upon accumulation. The intermediate variable *y(t)* is expected to correspond with concentration of plasticity related enzymes such as CaMKII or PP1 (Graupner and Brunel, 2007).

We first consider the effect of inhibitory input to synaptic plasticity at nearby excitatory spines. A recent experimental result revealed that, in medium spiny neuron, synaptic connections from cortical excitatory neurons typically show anti-Hebbian type STDP under pairwise stimulation protocol, but if GABA-A receptor is blocked, STDP time window flips to Hebbian (Paille et al., 2013) (points in Fig. 2A). The proposed model can explain this phenomenon in the following way. Let us first consider the case when the presynaptic excitatory input arrives before the postsynaptic spike. If the GABAergic input is blocked, presynaptic and postsynaptic spikes jointly cause a large membrane depolarization at the excitatory spine. Subsequently, the calcium concentration rises up above the LTP threshold (red line in Fig 2B upper-right), hence inducing LTP after repetitive stimulation (red line in Fig 2B lower-right). In contrast, if the GABAergic input arrives coincidentally with the presynaptic input, depolarization at the excitatory spine is attenuated by negative current influx though the inhibitory synapse. As a result, calcium concentration cannot go up beyond the LTP threshold although it is still high enough to eventually cause LTD (black lines in Fig 2B right). Similarly when the postsynaptic spike arrives to the spine before the presynaptic spike does, without any GABAergic input, the presynaptic spike causes slow decay in the level of calcium concentration that may induce LTD (red lines in Fig 2B left). On the contrary, if the GABAergic input is provided simultaneously with the presynaptic input, slow decay in the calcium concentration is blocked because the inhibitory input causes hyperpolarization of the membrane potential at the excitatory spine. As a result, LTP is more likely achieved (black lines in Fig. 2B left). Therefore, when a GABAergic input arrives in coincidence with a presynaptic excitatory input, the STDP time window changes its sign in both pre-post and post-pre regimes (lines in Fig. 2A).

GABAergic effect on excitatory synaptic plasticity is also observed in CA1 (Hayama et al., 2013). In this case, post-pre stimulation does not induce LTD unless GABA uncaging is conducted near the excitatory spine right before the postsynaptic spike arrives at the spine, whereas LTP is induced by pre-post stimulation regardless of GABA uncaging (blue and cyan points in Fig. 2C). The proposed model can also replicate these results. In pre-post stimulation, due to positive feedback through NMDA receptor, the membrane potential of the spine shows strong depolarization even if inhibitory current is delivered through GABA uncaging (solid and dotted blue lines in Fig. 2D upper-right). Thus, LTP is caused after repetitive stimulation (blue lines in Fig. 2D lower-right). By contrast, in post-pre protocol, LTP/LTD effects tend to cancel each other in the absence of GABAergic input, whereas LTD becomes dominant under the influence of GABAergic input (dotted and solid blue lines in Fig. 2D left, respectively).

In addition to inhibitory-to-excitatory effect, excitatory-to-excitatory (E-to-E) effect is also observed in case of CA1 (Hayama et al., 2013). If GABA uncaging is performed right before postsynaptic firing, LTD is also observed in neighboring excitatory spines (green point in Fig. 2C right). This E-to-E heterosynaptic effect is not observed in the absence of GABAergic input (green points in Fig 2C left). Correspondingly, in the model, excitatory current influx from a nearby synapse causes mild potentiation of calcium concentration in cooperation with inhibitory current influx, hence eventually induces LTD (green lines in Fig 2D left). Note that for this E-to-E effect, interaction at latter stage of synaptic plasticity may also play a dominant role (Hayama et al., 2013).

The obtained results are mostly robust against parameter change, as long as associated parameter values satisfy certain relationships (Fig. 3A). In addition, we found that the coefficient for heterosynaptic inhibitory effect should be larger for fitting the result from the striatum experiment, compared to CA1 (Fig. 3B top). This is consistent with strong inhibition observed in striatum (Mallet et al., 2005). We also found that for reproducing the result from the CA1 experiment, a high NMDA/AMPA ratio is crucial, while the striatum model is rather robust against it, as long as calcium influx/outflux is modulated by NMDA receptors (Fig. 3B bottom).

### Phase transitions underlying h-STDP

In the previous section, we introduced a biophysical model to establish its relevance to the corresponding biological processes and get insight into the underlying mechanism. However, not all components of the model are necessary to reproduce the observed properties of h-STDP. Here, we provide a simple analytically tractable model to investigate the generality of the proposed mechanism.

To this end, we simplify the model to the one in which calcium level at a spine is directly modulated by pre-, post-, and heterosynaptic activities as given below,

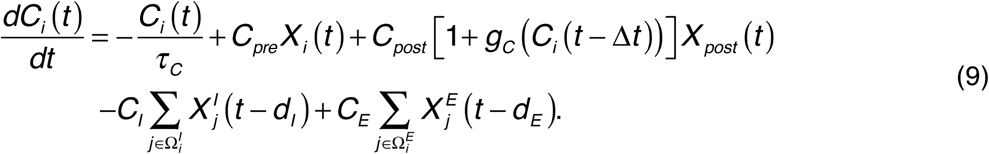

Here, *C_i_(t)* represents Ca^2+^ concentration at spine *i*, *X_i_* and *X_post_* represent presynaptic and postsynaptic spikes respectively, *d_I_* and *d_E_* are heterosynaptic delays, and 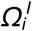 and 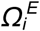 are the sets of neighboring inhibitory and excitatory synapses (see Reduced model in Methods for the details of the model). Despite simplicity, the model can qualitatively reproduce heterosynaptic effects observed in striatal and CA1 neurons, though the quantitative accuracy is degraded (Fig. 4A and B respectively). Importantly, the reduced model provides further analytical insights into the phenomena.

Let us consider how the inhibitory effect parameter *C_I_* controls I-to-E heterosynaptic effect observed in the CA1 experiment. If we characterize the shape of STDP time windows by the total number of its local minimum/maximum, the parameter space can be divided into several different phases (Fig. 4C). If LTP threshold *θ_p_* satisfies *C_pre_* < *θ_p_* < *C_post_*, Hebbian type STDP time window appears when the strength of heterosynaptic inhibitory effect *C_I_* satisfies (*C_post_* – *θ_p_*)*e*^*δ_I_*/*τ_C_*^ < *C_I_* < *C_pre_e*^*δ_I_*/*τ_C_*^ (upper orange-colored region in Fig. 4C; see Methods for the details of analysis). Here we defined *δ_I_* as the spike timing difference between inhibitory spike and presynaptic (postsynaptic) spikes in pre-post (post-pre) stimulation protocols. If *C_I_* is larger than *C_pre_*exp(*δ_I_*/*τ_C_*), a strong inhibitory effect causes LTD even in the pre-post regime (green-colored region in Fig. 4C), whereas LTD in the post-pre regime is suppressed when *C_I_* is smaller than (*C_pre_*-*θ_p_*)exp(*δ_I_*/*τ_C_*) (gray-colored region in Fig. 4C). Thus, heterosynaptic LTD observed in Figure 2C can be understood as the phase shift from the gray-colored region to the orange-colored region in Figure 4C, due to change in the inhibitory effect *C_I_*. This analysis further confirms that, for induction of heterosynaptic LTD, the heterosynaptic spike timing difference *δ_I_* should be smaller than the timescale of Ca^2+^ dynamics *τ_C_* (Hayama et al., 2013). This is because 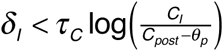 is necessary for a significant heterosynaptic LTD, and typically *C_I_* is smaller than *C_post_* and *θ_p_*. In addition, heterosynaptic suppression of pre-post LTP (green-colored region) is very unlikely to happen because *C_I_* > *C_pre_*exp(*δ_I_*/*τ_C_*) is necessary. This condition is difficult to satisfy even if *δ_I_*=0, because the heterosynaptic effect on Ca^2+^ dynamics in the spine is expected to be smaller than the homosynaptic effect (i.e. *C_I_* < *C_pre_*). A similar analysis is possible for E-to-E interaction although the phase diagram becomes complicated in this case (Fig. 4D; see Methods for details).

These analytical results revealed that the heterosynaptic effects are always observable if the parameters of calcium dynamics fall into a certain region in the parameter space, ensuring the robustness of h-STDP in our framework.

### h-STDP induces detailed dendritic E/I balance at dendritic hotspots

Results so far suggest that the proposed model gives a good approximation of h-STDP. We next study how this h-STDP rule shapes synaptic organization on the dendrite of a simulated neuron to investigate its possible functions. To this end, we first consider a model of a dendritic hotspot (Jia et al., 2010) that receives 10 excitatory inputs and one inhibitory input (Fig. 5A), because heterosynaptic effect is typically confined within 10*μ*m from the synapse (Hayama et al., 2013). Excitatory inputs are organized into 5 pairs, and each pair of excitatory synapses receives correlated inputs (Fig. 5B; see Dendritic hotspot model in Methods for details). In addition, the inhibitory input is correlated with one excitatory pair (in Fig. 5A, blue ones). Here, we assumed that postsynaptic activity follows a Poisson process with a fixed rate, because the influence of a single hotspot to the soma is usually negligible. In addition, we neglected the effect of morphology and hypothesized that heterosynaptic interaction occurs instantaneously within the hotspot. In this configuration, surprisingly, excitatory synapses correlated with the inhibitory input are potentiated while other synapses experience minor depression (Fig. 5C top). As a result, the dendritic membrane potential becomes less correlated with all hidden signals, because strong negative correlation with the blue signal is cancelled by potentiated excitatory inputs, while weak positive correlation with other signals are diminished due to LTD at corresponding excitatory synapses (Fig. 5C bottom). This GABA-driven potentiation is only observable when inhibitory activity is tightly correlated with excitatory activities, and becomes larger when inhibitory spike precedes excitatory spikes compared to the opposite case (Fig. 5D). In addition, we found that when heterosynaptic inhibitory effect γ_I_ is large enough to causes strong hyperpolarization at nearby synapses, depression is observed at correlated excitatory synapses (blue area in Fig. 5E) instead of potentiation (red area in Fig. 5E). However, as we saw in Figures 3 and 4, such a large inhibitory effect does not reproduce STDP experiments, thus unlikely to be observed in the actual brain. These results indicate that h-STDP induces dendrite-specific detailed E/I balance by potentiating excitatory synapses correlated with inhibitory synapses.

To reveal the underlying mechanism of this E/I balance generation, from the simulation data, we calculated the probability of calcium level being above the LTD/LTP thresholds after a presynaptic spike. The probability of LTP occurrence shows similar trajectories after a presynaptic spike, regardless of whether presynaptic activity is correlated with inhibitory input or not (blue and gray dotted lines in Fig. 5F, respectively). On the other hand, the maximum probability of LTD occurrence is significantly lower for spines correlated with inhibitory inputs (blue vs. gray solid line in Fig. 5F), although the probability goes up after the presynaptic spike in both cases. This asymmetry between LTP and LTD can be understood in the following way; LTD is mainly caused when the presynaptic neuron fires at a low firing rate and the postsynaptic neuron remains silent both in the experiment (Malenka and Bear, 2004) and in the model (gray line in Fig. 5G). However, if inhibitory input arrives at a nearby dendrite in coincidence, calcium boost caused by excitatory presynaptic input is attenuated by heterosynaptic inhibitory effect (black line in Fig. 5G). As a result, LTD is shunted by correlated inhibitory inputs. On the other hand, LTP is mainly caused by coincidence between pre and postsynaptic spikes, which induces a large increase in calcium level that overwhelms the attenuation by the heterosynaptic inhibitory effect. Thus, inhibitory activity at a nearby site does not prevent LTP at correlated excitatory synapses (Fig. 5H). Therefore, correlated spines experiences less depression, hence tend to be potentiated as a net sum.

To check the generality of the observed dendritic E/I balance, we extended the model to a two-layered single cell (Poirazi et al., 2003) by modeling each branch with one dendritic hotspot (Fig. 6A; see Two-layered neuron model in Methods for details), and investigated the dendritic organization by h-STDP. Here, we introduced 10 ms delay between excitatory and inhibitory stimulation(Froemke, 2015). Even in this case, when the postsynaptic neuron receives input from various neurons with different selectivity, each dendritic hotspot shapes its excitatory synaptic organization based on the selectivity of its inhibitory input (Fig. 6BC; the frame colors of Fig. 6B represent the inhibitory selectivities). As a result, excitatory synapses on the dendritic tree become clustered as observed in previous experiments (Kleindienst et al., 2011)(Takahashi et al., 2012). Note that, in our model, this clustering of excitatory synapses is caused by common inhibitory inputs instead of direct interaction among excitatory spines.

We further investigated the possible function of this synaptic organization in information processing. To this end, we consecutively presented the five stimuli to the two-layered neuron model (Fig. 6E). Before the learning, the neuron shows almost constant response to the stimulation with a small dip at the change points (Fig. 6E top). In contrast, after the learning, the neuron shows transient bursting activity immediately after the stimulus is changed to the next one, and rapidly returns to an almost silent state (Fig. 6E middle). Hence, by h-STDP, a neuron can acquire sensitivity toward abrupt changes in stimuli (Fig. 6D and 6E bottom).

### h-STDP explains critical period plasticity of binocular matching

Results so far indicate that h-STDP induces GABA-driven synaptic reorganization that enriches dendritic computation. To investigate its relationship with the developmental plasticity, we next consider a model of critical period plasticity in binocular matching (Wang et al., 2010)(Wang et al., 2013). In mice, one week after the eye opening, typically, binocular neurons in V1 still have different orientation selectivity for inputs from two eyes. Nevertheless, two more weeks after, selective orientations for both eyes get closer, and eventually they almost coincide with each other (Wang et al., 2010). Moreover, this binocular matching is disrupted by accelerating inhibitory maturation (Wang et al., 2013). Thus, expectedly, activity of inhibitory neurons play a crucial role in binocular matching in addition to Hebbian plasticity at excitatory synapses.

We modeled this process with a two-layered single cell model introduced in Figure 6 (Fig. 7A right; see Two-layered neuron model in Methods for details). Input spike trains were modeled as rate modulated Poisson processes driven by a circular variable *θ*, which corresponds to the direction of moving visual stimuli. We assumed followings: (i) inputs from ipsi-and contralateral eyes already have some weak orientation selectivity at the eye opening (Wang et al., 2010)(Espinosa and Stryker, 2012), (ii) Inhibitory cells are driven by both ipsi-and contralateral eyes (Yazaki-Sugiyama et al., 2009)(Kuhlman et al., 2011), (iii) The average selectivity of inhibitory inputs comes in between the selectivity for ipsilateral and contralateral excitatory inputs (Fig. 7A left). The last assumption has not yet been supported from experimental evidence, but if inhibition is provided from neighboring interneurons, these inhibitory neurons are likely to be driven by similar sets of feedforward excitatory inputs to those driving the output neuron. Here, we consider direction selectivity instead of orientation selectivity for mathematical convenience, but the same argument holds for the latter.

In the simulation, we first run the process without inhibition then introduced GABAergic inputs after a while (red lines in Fig. 7B,E represent the starting points of inhibitory inputs), because maturation of inhibitory neurons typically occurs in a later stage of the development (Hensch, 2005). Upon the introduction of inhibition, in each branch, the mean preferred direction of excitatory synapses converges to that of the local inhibition owing to heterosynaptic plasticity (Fig. 7B top; see Methods for details of evaluation methods), though synaptic weight development was biased toward the selectivity of the postsynaptic neuron (Fig. 7D; here, the bias is toward the right side). This dendritic E/I balancing shrinks the difference between ipsilateral and contralateral selectivity on average, because both of them get closer to the inhibitory selectivity (Fig. 7B middle). As a result, binocular selectivity becomes stronger (Fig. 7B bottom), and the responses for monocular inputs approximately coincide with each other (Fig. 7C right). Deprivation of contralateral inputs immediately after the introduction of inhibition blocks binocular matching (Fig. 7E), as expected from the experiment (Wang et al., 2010).

In addition, precocious GABA maturation is known to disrupt binocular matching (Wang et al., 2013). Our model suggests that the disruption is possibly related to the violation of the third assumption in the model. When the direction of the mean inhibitory selectivity is far different from both ipsilateral and contralateral selectivity (in Fig. 7F, at the parameter regions outside of the area surrounded by purple and green lines), h-STDP does not work effectively (Fig. 7F top), and the difference between ipsi-and contralateral inputs is not reduced (Fig. 7F middle). As a result, binocular direction selectivity is not improved by learning (Fig. 7F bottom). These results indicate that GABA-maturation and resultant h-STDP are an important part of the underlying mechanisms of critical period plasticity in binocular matching.

## Discussion

In this study, we first showed that a calcium-based plasticity model robustly captures several characteristics of plasticity-related interaction between neighboring synapses in millisecond timescale, by introducing heterosynaptic interaction terms (Fig. 2-4). Based on this proposed model, we next investigated the possible functions of h-STDP. Our study revealed that h-STDP causes the detailed dendritic E/I balance on dendritic hotspots (Fig. 5, 6), which is beneficial for change detection (Fig. 6). Furthermore, we found that h-STDP can induce binocular matching upon GABA maturation, and support an accurate input estimation (Fig. 7).

### Experimental predictions

Our study provides three experimental testable predictions: First, our results provide a hypothesis for synaptic organization on dendritic tree. It is known that excitatory synaptic inputs to a dendritic hotspot often show correlated activities (Kleindienst et al., 2011)(Takahashi et al., 2012). Our results indicate that an inhibitory input may also be correlated to excitatory inputs projecting to the nearby dendrite (Fig. 5,6), especially on a dendritic tree of an excitatory neuron that is sensitive to changes in the external environment (Fig. 6,7). Moreover, the model explains why feature selectivity of these spines only shows a weak similarity despite their correlations (Jia et al., 2010)(Chen et al., 2011). When a synaptic cluster is carved by the heterosynaptic effect of common inhibitory inputs, not by excitatory-to-excitatory interactions, variability of feature selectivity within the cluster tends to be large, because inhibitory neurons typically have a wider feature selectivity than excitatory neurons (Ma et al., 2010)(Moore and Wehr, 2013). In addition, it should also be noted that, E-to-E heterosynaptic LTP is typically induced as a meta-plasticity in the timescale of minutes (Harvey and Svoboda, 2007), which itself is not sufficient to create a correlation-based synaptic cluster.

Secondly, the results in Figure 5 indicate that LTD at an excitatory synapse is cancelled out by coincident inhibitory inputs to the nearby dendrite. Thus, LTD by low frequency stimuli (Malenka and Bear, 2004) can be attenuated by coincident GABA uncaging around the stimulated spine. Note that this result would not contradict with GABA-driven heterosynaptic LTD observed in paired stimulation, because in that experiment, the excitatory spine was presumably overexcited for inducing LTD in the absence of GABA (Hayama et al., 2013). Indeed, coincident GABAergic inputs may induce hetersosynaptic LTD if combined with presynaptic stimulation at a moderately high frequency that itself does not cause LTD (Blaise and Bronzino, 2003).

The third implication of the model is about binocular matching. Our model indicates that GABA-maturation plays a critical role in binocular matching, and proposes a candidate mechanism for disruption of binocular matching by precocious GABA maturation (Wang et al., 2013)(Fig. 7). However, the phenomenon can also be explained by Hebbian plasticity plus some kind of meta-plasticity. If binocular matching is purely induced by Hebbian plasticity not through heterosynaptic mechanism, selective orientation after the matching should depend solely on the initial selectivity for monocular inputs, assuming that selectivity of presynaptic neurons remains the same. Especially when the contralateral input is larger than the ipsilateral input, the resultant selectivity should approximately coincide with the original contralateral selectivity. On the other hand, if the proposed mechanism takes part in the development, the consequent selectivity should also be influenced by the mean selectivity of inhibitory input neurons. Thus, long-term imaging of monocular selectivity at binocular neurons in V1 would reveal whether a covariance-based rule is sufficient enough to explain the phenomena, or some other mechanisms including the proposed one also play a major role in the shift.

### Carrier of heterosynaptic interaction

Heterosynaptic plasticity has been observed in various spatial and temporal scales, and arguably underlying molecular mechanisms are different for different spatiotemporal scales (Nishiyama and Yasuda, 2015). In the case of milliseconds-order interaction, single-atomic ions are strong candidates, because poly-atomic ions such as IP_3_ are too big to move rapidly from spine to spine (Santamaria et al., 2006). Suppose that changes in Ca^2+^ concentration at an un-stimulated spine are crucial for heterosynaptic plasticity, Ca^2+^ influx/outflux from either intra or extracellular sources are necessary for induction of heterosynaptic plasticity. Because inhibitory synaptic inputs often change the local Ca^2+^ concentration in the dendritic branch (Müllner et al., 2015), intracellular spreading of Ca^2+^ may be a major source for Ca^2+^ changes in nearby un-stimulated spines. At the same time, because inhibitory inputs significantly modulate the membrane voltage of local dendrite (Gidon and Segev, 2012), a synaptic input should strongly drive Ca^2+^ influx/outflux through NMDA and VDCC from extracellular sources even at nearby un-stimulated spines. In addition, most of intracellular calcium-ions exist within calcium-buffer (Higley and Sabatini, 2012), and arguably they are also important for induction of synaptic plasticity. In our model, both current-based interaction (Spine model) and calcium-based interaction (Reduced model) replicate the experimental results (Fig. 2 and 4, respectively). Nevertheless, our analytical study suggest that the heterosynaptic Ca^2+^ change typically needs to be comparable with the homosynaptic change in order to cause significant heterosynaptic plasticity through calcium-based interaction (Fig. 4C, D). Thus, our study implies possible importance of current-based interaction and spine specific influx/outflux of extracellular Ca^2+^ for heterosynaptic plasticity.

Note that heterosynaptic interaction does not need to work in milliseconds order to interfere with STDP. For instance, E-to-E heterosynaptic LTD can be initiated by spreading of LTD-related molecules, not by messengers of neural activity (Hayama et al., 2013). In addition, for a shift in STDP time window, changes in the ratio between calcium influx through NMDA and the influx through VDCC possibly play a crucial role (Paille et al., 2013).

### Inhibitory cell types

Somatostatin positive (SOM^2+^) inhibitory neurons are typically projected to the apical dendrite, their IPSP curves is shorter than the timescales of NMDA or Ca^2+^ dynamics (Markram et al., 2004), and they often show strong feature selectivity compared to other inhibitory neuron types (Ma et al., 2010). Thus, this inhibitory cell type is the likely candidate for heterosynaptic STDP. However, our results do not exclude parvalbumin positive (PV^2+^) inhibitory neurons, which usually have projections to proximal dendrites, and also are typically fast spiking (Markram et al., 2004). In particular, h-STDP through PV^2+^ cell may play important roles in critical period plasticity (Takesian and Hensch, 2013).

### Related theoretical studies

Previous biophysical simulation studies revealed that synaptic plasticity at excitatory synapse critically depends on inhibitory inputs at nearby dendrite (Cutsuridis, 2011)(Bar-Ilan et al., 2013)(Jedlicka et al., 2015), but these studies did not reveal much of the functional roles of the heterosynaptic plasticity. On the other hands, network modeling studies found that heterosynaptic plasticity provides a homeostatic mechanism (Chen et al., 2013)(Zenke et al., 2015), but in these models, heterosynaptic plasticity was modeled as a global homeostatic plasticity without any branch specificity, and the advantage over other homeostatic mechanisms was unclear. In this study, by considering intermediate abstraction with analytical but biologically plausible models, we proposed candidate mechanisms for experimental results that have not been modeled before, and revealed potential functions of h-STDP in neural circuit formation.

## Acknowledgements

The authors thank to Dr. Laurent Venance for kindly providing the experimental data. This work was partly supported by JSPS Doctorial Fellowship DC2 (NH), CREST JST, and KAKENHI No15H04265 and No16H01289 (TF).

